# Wild grass isolates of *Magnaporthe* (Syn. *Pyricularia*) spp. from Germany can cause blast disease on cereal crops

**DOI:** 10.1101/2022.08.29.505667

**Authors:** A. Cristina Barragan, Sergio M. Latorre, Paul G. Mock, Adeline Harant, Joe Win, Angus Malmgren, Hernán A. Burbano, Sophien Kamoun, Thorsten Langner

**Affiliations:** The Sainsbury Laboratory, University of East Anglia, Norwich Research Park, Norwich, UK; Centre for Life’s Origins and Evolution, Department of Genetics, Evolution and Environment, University College London, UK

**Keywords:** Host jump, blast fungus, wild plants, cereal crops, pathogen migration, disease surveillance

## Abstract

Pathogens that cause destructive crop diseases often infect wild host plants. However, surveys of major plant pathogens tend to be skewed towards cultivated crops and often neglect the wild hosts. Here, we report an emerging disease threat generated by the blast fungus *Magnaporthe* (Syn. *Pyricularia*) spp. in central Europe. We found that this notoriously devastating plant pathogen infects the wild grasses foxtail millet (*Setaria* spp.) and crabgrass (*Digitaria* spp.) in south-western Germany, a region previously deemed unfavorable for blast disease. Using phenotypic characterization and genomic analyses, we determined that the observed disease symptoms are associated with the S*etaria* spp.-infecting lineage of *M. oryzae* and its sister species *Magnaporthe grisea*. We showed that *M. oryzae* isolates can infect barley and wheat, thus highlighting the risk of pathogen spread to crops. In addition, *M. oryzae* isolates which co-occur in natural populations display compatible mating types and variable candidate effector gene content, which may enhance the pathogen’s adaptive potential. Our findings stress the risk of blast fungus infections expanding into European cereal crops through migration and host jumps. This underlines the importance for pathogen surveillance not only on cultivated crops, but also on wild host plants.

**Author Summary:** Wild plant species are often overlooked during pathogen virulence surveys. However, many of the diseases we observe in crops are a consequence of host jumps from wild to cultivated plant species. This is reminiscent of zoonotic diseases, where host jumps from wild animals to humans result in new disease outbreaks. Here, we report that the notoriously devastating blast fungus *Magnaporthe* (Syn. *Pyricularia*) spp. occurs on wild grasses in south-western Germany. This region, which is at the center of the European cereal belt, has so far been viewed as having unfavorable climatic conditions for the blast disease. The newly identified blast fungus isolates have the capacity to infect wheat and barley cultivars, highlighting the risk of the disease spreading to staple cereal crops. In addition, there is potential for sexual recombination in local populations, which may increase the evolutionary potential of the fungus and might facilitate host jumps to cereal crops. Our findings emphasize the urgent need for surveillance of plant diseases on both wild hosts and crops.

## Introduction

Plant disease outbreaks are increasing at an alarming rate that is exacerbated by global trade and climate change, thereby threatening food security across the globe [1]. In addition, plant pathogens can jump from wild to cultivated host plants, resulting in new crop diseases [2]. This is similar to zoonotic diseases, where transmission from wild animal species to humans result in new disease outbreaks [3]. Although there are ongoing efforts to organize and increase global surveillance of plant diseases [4], these often neglect pathogens of wild hosts which have the potential to cause epidemics in crops [5]. As a consequence, our understanding of the distribution of potentially harmful pathogens reservoirs on wild hosts remains limited. This is particularly worrisome for crop pathogens with wide host ranges that can infect both cultivated crops and wild host plants.

One plant pathogen with a wide host range and a propensity to jump between hosts, is the ascomycete blast fungus *Magnaporthe oryzae* (Syn. *Pyricularia oryzae*). *M. oryzae* infects over fifty wild and cultivated grass species, including staple cereal crops such as rice (*Oryza sativa*), wheat (*Triticum aestivum*) and barley (*Hordeum vulgare*) [6]. In contrast, its sister species *Magnaporthe grisea*, is mainly associated with crabgrass (*Digitaria* spp.) infections [7,8]. *M. oryzae* is thought to be undergoing incipient speciation into genetically distinct lineages, however, some inter-lineage genetic exchange does occur [9]. To date, at least ten *M. oryzae* lineages predominantly associated with a single grass host genus have been described [10]. *M. oryzae* is thought to have undergone host jumps multiple times from wild to cultivated grass species. The rice-infecting lineage, for example, likely originated after a host shift from the *M. oryzae* lineage infecting foxtail millet (*Setaria* spp.) [11], wheat blast emerged after a host jump from the *M. oryzae* lineage infecting ryegrass (*Lolium* spp.) [12], and a maize-infecting lineage has evolved after a host jump from barnyard grass (*Echinochloa crus-galli*) [13].

Blast fungus infections, especially on rice and wheat crops, have had dire economic and societal consequences. Wheat blast, for example, emerged in the mid 1980s and was initially restricted to South America, but has spread to South Asia (Bangladesh, 2016) and Africa (Zambia, 2018) in recent years, resulting in annual yield losses reaching up to 50% [14–16]. In Europe, blast outbreaks are largely restricted to southern countries due to more favorable environmental conditions for the fungus, which is especially well-documented for rice blast in the Mediterranean [17]. A phytosanitary risk assessment by the Julius Kühn Institute and the European and Mediterranean Plant Protection Organization (EPPO), last updated in 2019, stated “So far, the fungus *M. oryzae* is not present in Germany but it already occurs in other EU-Member States. (…) Presumably, *M. oryzae* will not be able to establish outdoors in Germany due to unsuitable climate conditions” (EPPO; 2019). So far, this major cereal producing region at the center of the European cereal belt has been spared from blast disease.

Climate change will likely result in newly emerging crop diseases across the European continent and could result in environmental conditions suitable for blast disease. Average near-surface temperatures in Europe have already increased by 1.9°C compared to pre-industrial levels (EEA; 2022), with temperatures expected to continue to rise in the next decades [18]. Climate change has resulted in shifts in the geographical range of crop pests and pathogens [19] which have been documented in a myriad of fungal pathogens [20]. For example, the *Fusarium graminearum* species complex, which causes Fusarium head blight disease in multiple cereal crops, is typically present in warm, humid conditions. However, in recent years it has become more common in Germany, where it poses a threat to maize production, likely enabled by increasing temperatures [21].

We surveyed grass species for blast disease symptoms in the summer of 2021 to better understand the diversity and distribution of the blast fungus in southern and central Europe. We observed typical symptoms of blast fungus infections on the wild host plants *Setaria* spp. and *Digitaria* spp. in south-western Germany, a region located in the center of the European cereal belt. Using phenotypic characterization and whole genome analyses, we placed the collected isolates in the general *Magnaporthe* phylogenetic framework and determined that the observed disease symptoms are associated with the *Setaria* spp.-infecting lineage of *M. oryzae* and its sister species *M. grisea*, which infects *Digitaria* spp. We found that the *M. oryzae* isolates can infect susceptible wheat (*Triticum aestivum*) and barley (*Hordeum vulgare*) cultivars under laboratory conditions. In addition, some co-occurring isolates carry opposite mating types and display virulence effector diversity, potentially increasing the adaptive capacity of the fungus. Together, our findings highlight the risk that climate conditions in central Europe may become more conductive for blast diseases. Furthermore, blast fungus lineages infecting wild grasses may adapt to locally cultivated cereal crops, posing a risk to key agricultural sectors in the region. Ultimately, our results point towards a need for increased pathogen surveillance, not only on crops, but also on wild grass hosts.

## Results

### Disease symptoms and morphological features suggest the presence of the blast fungus in Germany

We observed plants which displayed elliptical lesions with a necrotic center and dark edges, typical for blast disease on *Setaria* spp. (**Fig 1A**) and *Digitaria* spp. plants (**Fig 1B**) in south-western Germany in August 2021 (**Fig 1C, 1D** and **Table S1**). We collected aerial material of representative individuals from infected plants and transported them to the laboratory for further analysis. After processing of infected plant material, we cultured 18 single spore isolates from nine infected plant samples: 16 isolates from infected *Setaria* spp. and two isolates from infected *Digitaria* spp. While the colony morphology of the two *Digitaria* spp.-infecting isolates was similar, *Setaria* spp.-infecting isolates displayed diverse colony morphologies. Eight of these isolates formed entirely white, fluffy colonies, whereas eight developed a gray center indicative of sporulation or high pigmentation. We selected two *Setaria* spp.-infecting isolates with contrasting colony morphologies (GE10A2 and GE12B) and the two *Digitaria* spp.-infecting isolates (GE3 and GE16_2) for further investigation (**Fig 1E** and **1F**). Microscopic characterization confirmed typical blast fungus morphology, with two septate, spindle-shaped spores, that formed characteristic single-celled appressoria on the hydrophobic surface of cover slips (**Fig 1G** and **1H**). Taken together, our results show that disease symptoms, colony, spore and appressorium morphology of the samples we collected in Germany are consistent with the blast fungus.

**Fig 1.**
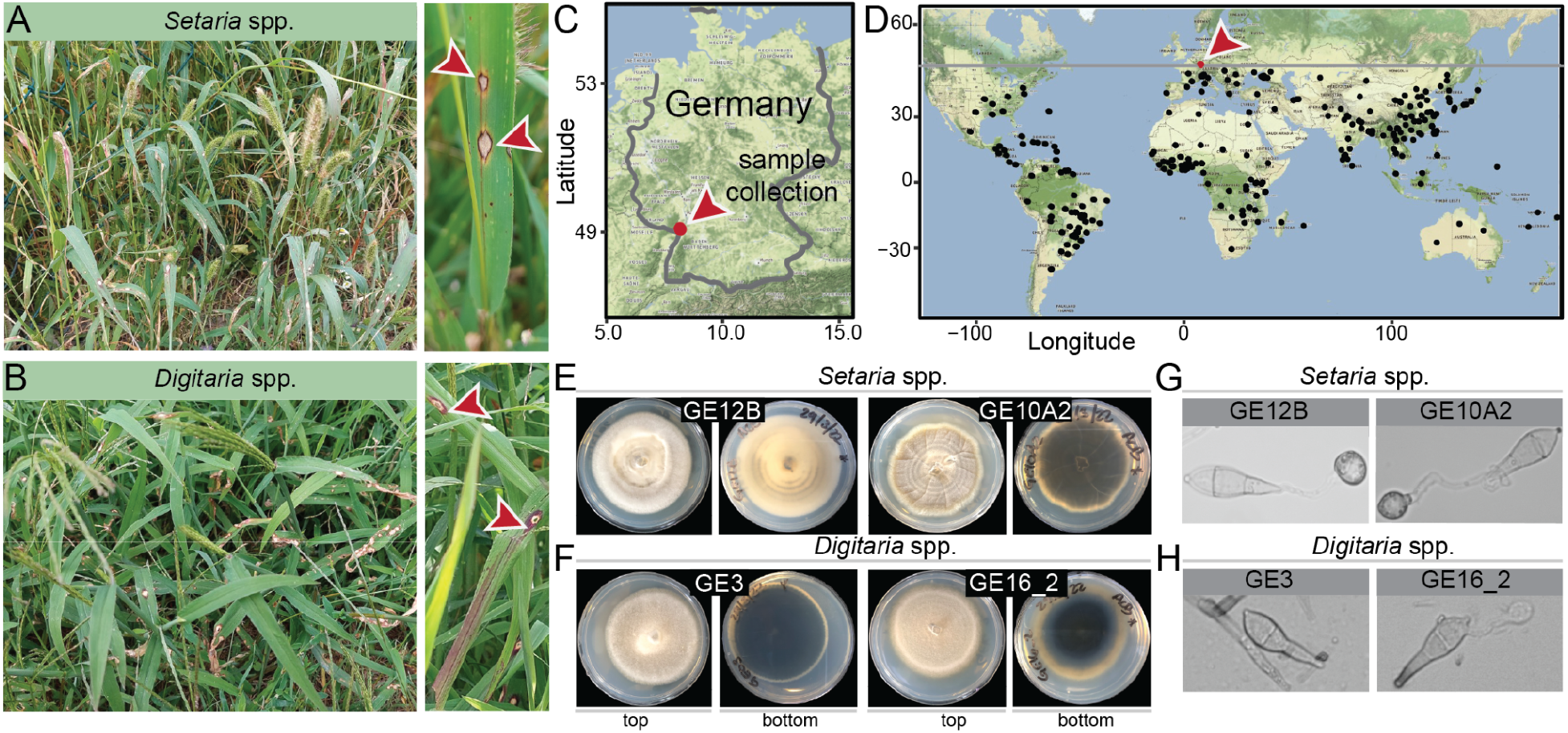
Disease symptoms and pathogen morphology suggest the presence of the blast fungus in Germany. **A-B**. Characteristic blast fungus lesions on *Setaria* spp. (A) and *Digitaria* spp. (B) observed in south-western Germany. Red arrows indicate lesions. **C**. Location of infected German samples (red). **D**. Locations of blast fungus occurrences (see Methods, Table S1), and collected plant samples in south-western Germany (red). Latitude of the sampling site in Germany is indicated as a gray line. **E-F**. Mycelium morphology of representative isolates infecting *Setaria* spp. (E) and *Digitaria* spp. (F). **G-H**. Spore and appressorium morphology from the same isolates as in E-F.

### Whole genome analysis confirms presence of the blast fungus *M. oryzae* and its sister species *M. grisea* in Germany

To define the precise identity of the isolated fungi, we selected the four isolates described above for whole genome sequencing using long and short read sequencing technologies [22]. This resulted in four high quality genome assemblies (13-29 contigs) with a BUSCO completeness score 98.8-98.9% (**Table 1**).

**Table 1.** Summary statistics of polished genome assemblies of German *Magnaporthe* isolates.

| Sample name | Assembly length (bp) | No. contigs | Contig N50 (bp) | Largest contig (bp) | Mean coverage (x fold) | Complete BUSCO<sup>1</sup> | GenBank Accession |
| --- | --- | --- | --- | --- | --- | --- | --- |
| GE12B | 44,287,786 | 20 | 5,065,357 | 8,374,180 | 109 | 98.90% | GCA_944989255.1 |
| GE10A2 | 44,757,806 | 29 | 6,972,091 | 11,001,122 | 123 | 98.90% | GCA_944989335.1 |
| GE3 | 43,441,267 | 15 | 5,238,139 | 8,063,860 | 105 | 98.80% | GCA_944989265.1 |
| GE16_2 | 43,662,581 | 13 | 6,231,040 | 8,069,839 | 146 | 98.80% | GCA_944952825.1 |

<sup>1</sup>BUSCO=Benchmarking Universal Single-Copy Orthologs. BUSCO was run using ascomycota\_odb10 dataset.

Next, we mapped short reads of these four isolates, in addition to 413 publicly available *M. grisea* and *M. oryzae* samples (**Table S2**), to the reference assembly MG08 (isolate 70-15) [23]. We then performed variant calling and determined their genetic relationship by creating a genome-wide SNP-based Neighbor-Joining (NJ) tree (**Fig S1A**). For easier visualization, we subsampled 80 isolates representative for *M. grisea* and the different *M. oryzae* lineages (**Fig 2**). In both cases, the German *Setaria* spp.-infecting samples clustered with other members of the *Setaria* spp.-infecting *M. oryzae* lineage, whereas the *Digitaria* spp.-infecting samples clustered with Dig41, a *Digitaria* spp.-infecting *M. grisea* isolate typically used as an outgroup in *M. oryzae* phylogenies [24,25]. The German blast fungus isolates were most similar to each other. For the *Setaria* spp.-infecting *M. oryzae*, this was confirmed by creating a SNP-based NJ-tree for this lineage only (**Fig S1B**).

**Fig 2.**
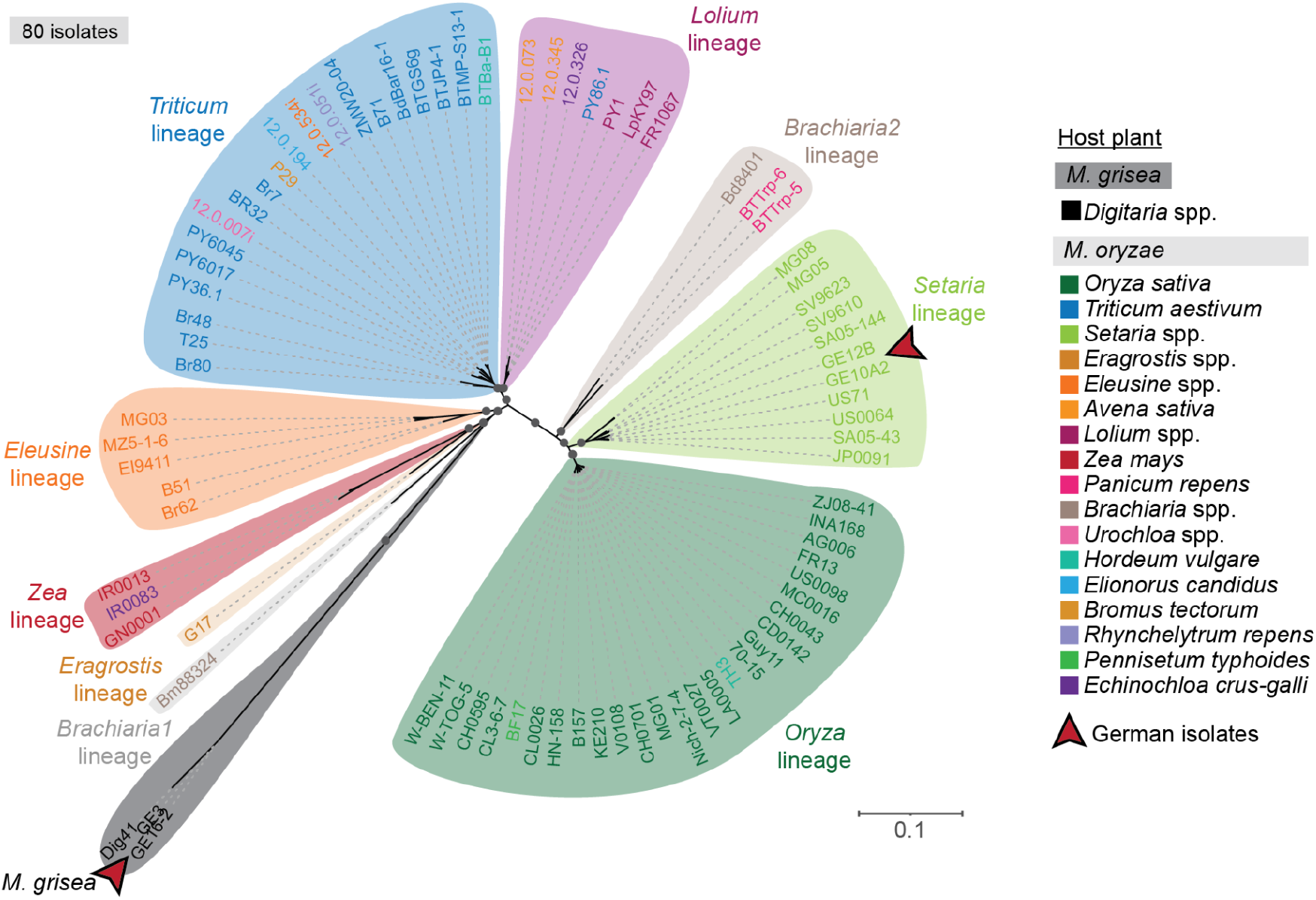
*M. oryzae* and its sister species *M. grisea* are present in Germany. Genome-wide, SNP-based Neighbor-Joining (NJ) tree of 80 *Magnaporthe* isolates color-coded by their original host plant and genetic lineage. Red arrows highlight the German isolates. The *Digitaria* spp.-infecting isolates (GE3 and GE16_2) cluster with the *M. grisea* isolate Dig41 (black), while the *Setaria* spp.-infecting isolates (GE12B and GE10A2) cluster with isolates of the *Setaria* spp.*-*infecting *M. oryzae* lineage (light green). Scale bar indicates nucleotide substitutions per position. Relevant bootstrapping values ≥ 0.9 are shown as gray circles.

To avoid a potential reference-bias, and complementary to the genome-wide SNP-based NJ trees, we used unmapped sequencing reads to estimate k-mer-based pairwise distances between the same subset of 80 representative *M. oryzae* and *M*.*grisea* isolates used to create NJ trees (**Table S2**). The relationship between all isolates and their grouping into host-specific lineages was consistent in both analyses and corroborated the clustering of the German isolates based on genome-wide SNPs (**Fig S2A**). We repeated the reference-free, k-mer based classification using increased stringency (see Methods) which confirmed robustness of this method and the identity of German blast fungus isolates (**Fig S2B**). In summary, our genomic analysis confirmed the presence of two blast fungus species, *M. oryzae* and *M. grisea*, in Germany and highlights the robustness of the k-mer based, reference free method for pathogen identification.

### German *M. oryzae* isolates can infect wheat and barley cultivars

Next, we investigated whether the blast fungus isolates we collected from wild grasses have the capacity to infect cereal crops, such as barley and wheat, which are extensively cultivated in central Europe. We performed leaf drop infection assays using the four sequenced isolates on the barley cultivars Nigrate and Golden Promise, and the wheat cultivars Fielder and Chinese Spring. As a positive control, we used the wheat-infecting blast fungus isolate BTMP-S13-1, which belongs to the highly virulent pandemic clonal wheat blast lineage (“B71 lineage”) that caused major outbreaks after its introduction to Bangladesh in 2016 [14,15,26–29]. The four tested German isolates showed differential virulence in these cultivars under laboratory conditions. While *M. grisea* isolates were largely avirulent on these plants, the *Setaria* spp.-infecting *M. oryzae* isolates caused disease symptoms in the form of progressing lesion development (**Fig 3**). Disease progression was most pronounced in the susceptible barley cultivars Nigrate and Golden Promise, but progressing lesions also formed occasionally on the wheat cultivars Fielder and Chinese Spring.

**Fig 3.**
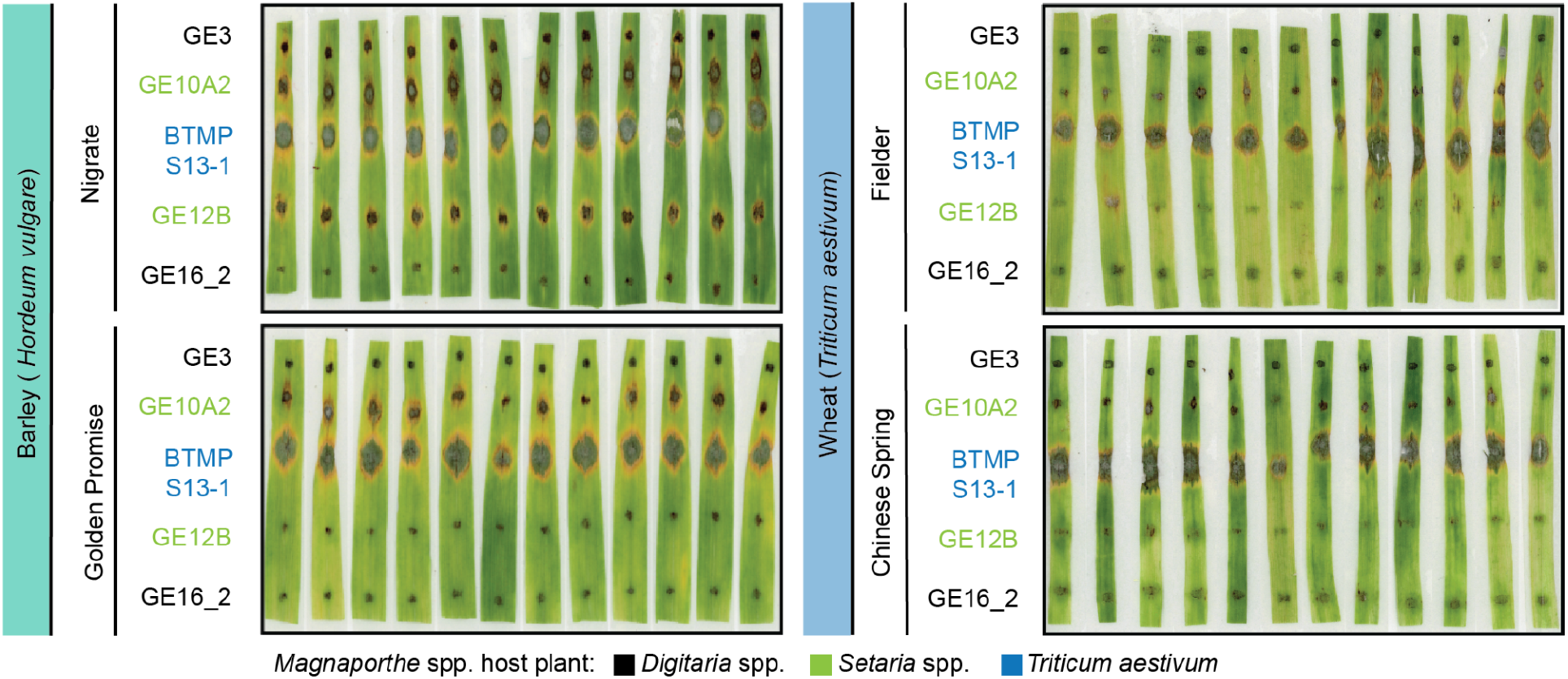
German *M. oryzae* isolates can infect wheat and barley cultivars under laboratory conditions. Detached leaf virulence assay of 10-14 day old barley cultivars Golden Promise and Nigrate, and wheat cultivars Fielder and Chinese Spring. Among the German blast fungus isolates, lesion development was most pronounced for the *Setaria* spp.-infecting *M. oryzae* isolates (light green) on susceptible barley cultivars. Disease phenotypes were scored 6 days after infection.

To test whether the *Setaria* spp.-infecting *M. oryzae* isolates, which cause visible lesion development, can complete their infection cycle on barley and wheat, we transferred the lesions from infection assays to 2% water agar to induce sporulation for 24 hours. We observed formation of conidiophores on all four tested host cultivars from infections with the *M. oryzae* isolate GE12B, and from infections with isolate GE10A2 on the barley cultivar Nigrate. To determine whether the spores that are produced on these plants are viable, we transferred single spores to CM plates and observed colony formation (**Fig S3**). This suggests these isolates can complete their asexual cycle on wheat and barley cultivars under laboratory conditions. Taken together, our results suggest that the *M. oryzae* population present in Germany has the potential to cause disease on cereal crops.

### Mating types and effector candidate genes vary across German blast fungus isolates

Sexual mating in the blast fungus requires the co-occurrence of compatible mating types [30]. To determine the potential for sexual reproduction in local blast fungus populations in Germany, we analyzed the mating types of the isolates we collected. We used a BLASTN sequence similarity-based approach using the previously described MAT1-1 and MAT1-2 mating type genes [31,32]. Both *M. grisea* isolates GE3 and GE16_2, and the *Setaria* spp.-infecting *M. oryzae* strain GE12B, carry the MAT1-1 mating type locus idiomorph, while the *Setaria* spp.*-*infecting strain GE10A2 carries the MAT1-2 idiomorph. This demonstrates that *M. oryzae* isolates of opposite mating types co-occur in local populations, highlighting the potential of hybridization between isolates.

The ability of the blast fungus to cause disease and undergo host jumps, is largely determined by its effector gene content which can vary even among closely related isolates, resulting in differential virulence to host plants [12,24,33,34]. We therefore analyzed presence/absence variation of candidate effector genes in the German blast fungus isolates, and twelve publicly available *M. oryzae* and *M. grisea* isolates isolated from *Setaria* spp. or *Digitaria* spp. based on sequence similarity to 178 candidate effectors which showed sequence similarity to validated effectors, or were predicted MAX (*Magnaporthe* avirulence [Avr] and ToxB like) effectors [35,36]. The latter are sequence-unrelated but structurally conserved fungal effectors that are expanded in *Magnaporthe*, which represent half of experimentally validated effectors [37] and comprise two-thirds of the 178 effectors candidates tested.

From 178 candidate effector sequences, 165 (93%) showed presence/absence variation across the sixteen *Magnaporthe* isolates compared. Candidate effector content was most similar among isolates that infect the same host species, in agreement with previous observations [38] (**Fig 4**). Notably, among the German isolates, six predicted effectors, including three predicted and one validated MAX effector, showed presence/absence variation between the two *M. oryzae* isolates infecting *Setaria* spp., GE10A2 and GE12B. In the *Digitaria* spp.-infecting isolates GE3 and GE16_2, two predicted effectors, including a predicted MAX effector, displayed presence/absence variation (**Table S3**). These variable effectors included the well-characterized AVR-Pita and AVR-Pia, which are associated with host-specialization in *M. oryzae* [39,40]. In summary, candidate effector content varies even between the genetically similar German blast fungus isolates.

**Fig 4.**
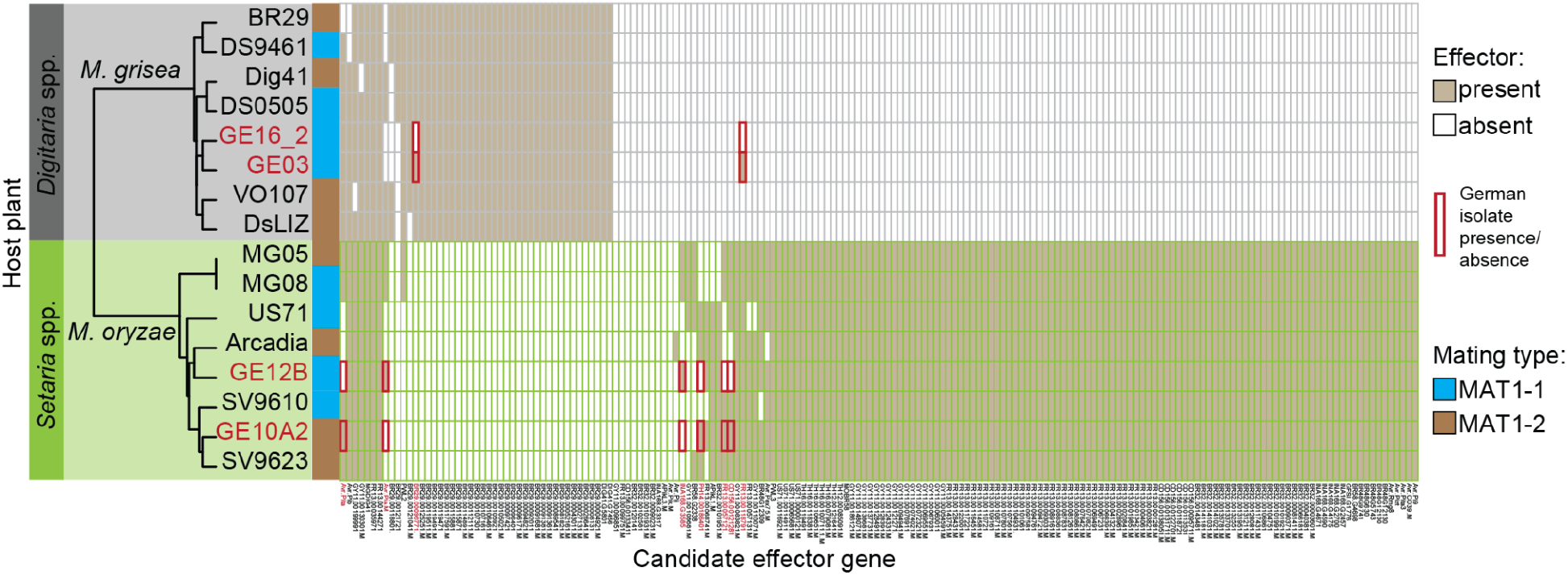
Opposite mating types and variable candidate effector repertoires are present in the German blast fungus. Mating types of each isolate is shown in blue (MAT1-1) and brown (MAT1-2). Co-occurring German blast fungus isolates infecting *Setaria* spp. carry opposite mating types. Presence/absence variation of effector candidates in isolates infecting *Setaria* spp. (light green) and isolates infecting *Digitaria* spp. (gray). German isolate IDs are highlighted in red. Effector candidate genes which show presence/absence variation between the German samples infecting the same host are also highlighted in red. Dendrogram reflects similarity in candidate effector repertoire.

## Discussion

### Blast diseases may pose an emerging threat to central European cereal production

Blast diseases in Europe have so far mainly been observed in the Mediterranean and southern European regions ([CABI; 2022](https://www.cabi.org/isc/)). In contrast, environmental conditions in central and northern Europe are thought to be largely unfavorable for the blast fungus, and thus, blast disease has not been considered a major risk in Germany ([EPPO; 2019](https://pra.eppo.int/pra/5582c4e9-6c10-4f91-b132-e82c335c5fe0)). However, our findings show that the blast fungus *M. oryzae* and its sister species *M. grisea* can establish infections under natural conditions on wild grasses in Germany, pointing to the risk of cereal blast emerging as a crop disease in central Europe. Cereals dominate agriculture in the European Union and approximately one third of the Utilized Agricultural Area is occupied by cereals and another third by permanent grassland ([EUROSTAT; 2015](https://ec.europa.eu/eurostat/statistics-explained/index.php?title=Archive:Main_annual_crop_statistics)). In our laboratory experiments, German blast fungus isolates could infect wheat and barley, two major crops of the European cereal belt which includes northern France and the entire territory of Germany [41]. These two countries alone account for ∼39% of the wheat and ∼36% of barley production in the European Union ([USDA; 2014](https://ipad.fas.usda.gov/rssiws/al/europe_cropprod.aspx)).

### What is the risk of host jumps of the blast fungus in Central Europe?

Based on historical precedence and our observation that *Setaria* spp.*-*infecting *M. oryzae* isolates can cause disease on cereal crops, a host jump of the *M. oryzae* lineage infecting *Setaria* spp. to cultivated grass hosts is a latent threat in Central Europe. Similar to zoonotic diseases, where outbreaks of human diseases can often be traced back to wild animal infections [3], plant pathogens occasionally undergo host jumps or host range expansions from wild to cultivated plant host species followed by adaptation to the new host [42].

We show that *Setaria* spp.-infecting *M. oryzae* isolates can complete their infection cycle and reproduce asexually on barley and wheat under laboratory conditions. Although the significance of cross-infections of alternative hosts under natural conditions remains to be investigated, the risk of a host jump from these wild grasses to cereal crops in central Europe should not be ignored. In the past, the blast fungus has undergone jumps between wild and cultivated host plants on multiple occasions [11–13,43]. In addition, host cross-infectivity in the field appears to be relatively common, where a single blast fungus lineage infects multiple host species [25]. Examples are the rice blast fungus isolate TH3 which was collected from barley in Thailand, or the wheat blast fungus isolate P29, which was collected from cheatgrass (*Bromus tectorum*) in Paraguay (**Fig 2** and **Table S2**). Intermediate hosts, such as barley cultivars that are susceptible to most blast fungus lineages [44,45], may serve as “springboards” for rapid pathogen diversification under weak host selective pressure. This in turn may facilitate adaptation of the blast fungus to local cereal cultivars.

### Mating type and effector gene variation may increase the adaptive potential of the blast fungus in Europe

The potential of plant pathogens to adapt to changing environmental conditions and novel host plants may be exacerbated if sexual recombination occurs in populations where fertile isolates of opposite mating types co-occur. Our analysis shows that opposite mating types co-occur in a local *M. oryzae* population infecting *Setaria* spp. in Germany (**Fig 4**). Sexual recombination between fertile blast fungus isolates is thought to occur in nature and can readily be achieved under laboratory conditions [9,28,36,46]. In addition, genomic analyses suggest that recombination occurs in some lineages, although pandemic clonal lineages of rice and wheat blast tend to dominate isolates obtained from these crops [9,25,28,29,36,47]. We recently showed that an isolate of the pandemic clonal wheat blast lineage (“B71 lineage”) from Zambia can mate with an African finger miller isolate [28], further raising the possibility that recombination and hybridization between host-specialized lineages can drive the pathogen’s evolutionary potential.

We observed presence/absence variation of predicted candidate effector genes across the four German blast fungus isolates we characterized. Differential virulence and adaptation to alternative host plants largely depends on the pathogen effector gene content and the immune receptor (resistance gene, or short, *R*-gene) repertoire of the host population [24,38–40,48]. Two of the effector candidates that show presence/absence variation in German *Setaria* spp.-infecting isolates are the well-characterized avirulence effectors AVR-Pita and AVR-Pia, which are recognized by host plants that carry the corresponding immune receptors Pi-ta and Pia, respectively [39,40]. Although the genotype and *R*-gene content of German *Setaria* spp. hosts remain unknown, it is possible that *R*-gene diversity in these wild populations led to the observed variation of pathogen effector genes. Differences in blast disease susceptibility has been described for both cultivated (*Setaria italica*) and wild (*Setaria viridis*) *Setaria* spp. accessions [49] and homologs of several blast fungus resistance genes, including Pia (syn. RGA4/RGA5) have been identified in *Setaria* spp. genomes [50,51]. In summary, our data suggests that the presence of compatible mating types and standing genetic variation, most notably of candidate effector gene content, might contribute to the adaptive potential of blast fungus infecting wild grass populations in Germany.

### Climate change may promote a northward migration of the blast fungus in Europe

Climate conditions in central and northern Europe are generally viewed as unfavorable to support blast disease outbreaks. Prior to this report and based on our literature survey, blast disease has been reported as far north as Ukraine, Hungary, southern France, and northern Italy on the host plants rice (*Oryza sativa*) or ryegrass (*Lolium* spp.) (**Table S2**). In this report, we describe the northernmost occurrence of blast disease in Europe. Our finding that the blast fungus can cause disease in Germany under central European climate conditions should be of general interest to the plant pathology community and policy makers as it may point towards a potential northward migration of a highly devastating plant pathogen. While surveillance for blast disease in Germany is limited, historical climate data and future predictions [52–55] make it conceivable that, as temperatures rise, the blast fungus will spread further north into major cereal producing regions in Europe in the future [56]. How far north the blast fungus is established in Europe beyond the area we surveyed is unknown. This lack of knowledge is further exacerbated by the paucity of surveys on wild host species. We therefore urge the community to increase disease surveillance not only on crops, but also on wild hosts, as is routinely performed for zoonotic human diseases.

## Conclusion

The climatic conditions in Germany and the wider central European cereal belt have so far been considered unfavorable for proliferation and spread of the blast fungus. It is unclear how widespread the fungus is in this region, especially since occurrences of the blast fungus on uncultivated wild plants often remain undetected. The sample location reported here is in the center of the southern-German Upper Rhine valley, one of the warmest areas in Germany [57]. It is thus possible that the spread of the blast disease is limited to this geographical region. Nevertheless, our study exposes the blast fungus as an emerging threat to cereal crops grown in central Europe and highlights the importance for plant pathogen surveillance not just on crop plants but also on wild hosts. This is increasingly important in the face of changing pathogen dynamics as a consequence of climate change, and shifting global trade routes propelled by international conflicts.

## Materials and Methods

### Sample collection and phenotypic pathogen characterization

In mid-August of 2021, we observed blast disease symptoms on the Poaceae hosts wild foxtail millet (*Setaria* spp.) and crabgrass (*Digitaria* spp.) in the state of Rhineland-Palatinate, in south-western Germany (Latitude 49.08314° N, Longitude 8.20417° E). Infected plants showed elliptical, gray, necrotic lesions with dark edges, typical for *M. oryzae* and *M. grisea*, with lesions in *Digitaria* spp. additionally displaying a characteristic purple tint on lesion edges. We collected aerial parts of diseased plant material and air dried them for further inspection in the laboratory. To induce sporulation, we placed the infected leaves into a moist chamber containing 2% water agar for 24-48 hours at room temperature until two-septate conidia formed. We then transferred sporulating lesions to complete medium (CM) for 7-14 days in a growth chamber at 24°C with a 12 hour light period to induce growth of mycelium and sporulation. Spores were harvested in sterile water and plated on 2% water agar for single spore isolation. After 24 hours, germinated single spores were transferred to CM plates and grown for 8-12 days in a growth chamber at 24°C, which led to the formation of typical mycelial colonies. We observed conidiospores of representative samples microscopically and confirmed spore morphology and formation of appressoria in 50 µl sterile water on hydrophobic coverslips (Menzel-Gläser; 22 × 50 mm #1, Part. No. 15787582).

### Visualization of worldwide blast fungus distribution

Maps were created with the R-package ggmap (v3.0) [58]. Locations of blast fungus samples plotted are those of the 417 *M. oryzae* and *M. grisea* samples which were used for SNP-based phylogenetic analyses (**Table S2**). In addition, locations of blast fungus reports from Russia [59] and Korea [60], as well as the location of reports of rice and wheat blast fungus from CABI, the Invasive Species Compendium, were plotted. If the exact coordinates of a sample were unknown, the coordinates of the county’s geographical center were plotted (**Table S1**).

### Pathogen infection assays

The third leaf of 10-14 day old seedlings (three leaves stage) of wheat cultivars Fielder and Chinese Spring, and barley cultivars Golden Promise and Nigrate were drop inoculated with 10 µl spore suspension (1×10^5^ spores ml^-1^) of two *M. oryzae* single-spore isolates infecting *Setaria* spp. (strain ID: GE12B and GE10A2) and two *M. grisea* single spore isolates infecting *Digitaria* spp. (strain ID: GE3 and GE16_2), in addition to a pandemic wheat blast strain originating from Bangladesh, BTMP-S13-1, as a positive control. Twelve leaves per host genotype were tested for virulence of the isolates. The inoculated detached leaves were incubated in a growth room at 24°C with a 12 hour light period. Disease phenotypes were scored 6 days after drop infection. To assess whether *M. oryzae* isolates can complete their infection cycle and proliferate in infected host tissue, we reisolated spores from infected lesions by placing lesions on 2% water agar plates at room temperature overnight to induce sporulation. We confirmed generation of conidiophores on the plant surface microscopically before transferring single spores to CM plates for proliferation at 24°C. Viable spores were isolated from *M. oryzae* infections, suggesting these isolates can cross-infect and proliferate on susceptible barley and wheat cultivars under laboratory conditions.

### Genome sequencing and assembly

We whole-genome sequenced two representative single spore isolates infecting *Setaria* spp. (isolate ID GE12B and GE10A2) and two infecting *Digitaria* spp. (isolate ID GE3 and GE16_2) as described in [22]. For this purpose, we extracted high molecular weight DNA following the protocol described in [59]. Sequencing runs were performed by Future Genomics Technologies (Leiden, The Netherlands) using the PromethION sequencing platform (Oxford Nanopore Technologies, Oxford, UK). All samples were sequenced with a depth between 105X and 146X and a mean read length of >20kb (read N50 >30kb) (**Tables S4** and **S5**). Long reads were assembled into contigs and corrected using Flye (v2.9-b1768) [60], and polished with long reads using Medaka (v1.4.3) (https://github.com/nanoporetech/medaka). All assemblies were additionally polished using short read Illumina sequencing technologies (San Diego, USA) through two consecutive iterations of Pilon (v1.23) [61]. The resulting assemblies were both of high quality and contiguity, with a BUSCO [62] completeness score of 98.8-98.9% (ascomycota_odb10 database) and 13-29 contigs (**Table 1** and **S6**). Sequencing reads were deposited in the European Nucleotide Archive (ENA) under study accession number PRJEB53658 and run accessions ERR9866173 (GE12B), ERR9866254 (GE10A2), ERR9883755 and ERR9883756 (GE3) and ERR9866258 and ERR9866259 (GE16_2). In addition, the four whole-genome assemblies are available under GenBank accession numbers GCA_944989255.1 (GE12B), GCA_944989335.1 (GE10A2), GCA_944989265.1 (GE3) and GCA_944952825.1 (GE16_2).

### Mapping, variant calling and genetic clustering of blast fungus isolates

Illumina short reads of the four German blast fungus isolates sequenced and 413 *M. oryzae* and *M. grisea* isolate infecting different host plants (**Table S2**) were trimmed using AdapterRemoval (v2.3.1) [61] and then mapped to the 70-15 *M. oryzae* reference genome (MG08) using bwa-mem (v0.7.17) [62] with default parameters (https://github.com/smlatorreo/Magnaporthe-Germany). Average read depths were estimated using the samtools depth command [63] (**Table S3**). Variant identification was performed using GATK (v4.1.4.0) [64]. High-quality SNPs were filtered based on the Quality-by-Depth (QD) parameter using GATK’s VariantFiltration. Only SNPs within one standard deviation of the median value of QD scores across all SNPs were kept [65]. We kept only biallelic sites, and a total of 3’035,335 SNPs were obtained. For analyses across all 417 samples, only positions with no missing data (--max-missing 1.0) were kept using VCFtools (v0.1.14), which resulted in 441,820 SNPs. SNPs displaying high levels of linkage disequilibrium were then pruned using PLINK1.9 (--indep-pairwise 50 10 0.5), which resulted in 59,945 SNPs. These SNPs were then used for the construction of Neighbor-Joining (NJ) trees using MEGA (v10.2.4) [66], with 150 bootstraps which were visualized in iTol [67] (https://itol.embl.de/tree/14915519290215291654101261). For easier visualization, the same process was repeated for 80 representative *Magnaporthe* spp. isolates (**Table S2**). Here, the --max-missing 1.0 filter resulted in 1’066,004 SNPs and --indep-pairwise 50 10 0.5 filtering resulted in 94,637 SNPs (https://itol.embl.de/tree/14915519290129221654184627). In order to confirm the phylogenetic relationship between *Setaria* spp.-infecting *M. oryzae*, this filtering process was repeated with members of this lineage only (--max-missing 1.0 filtering resulted in 1’835,862 SNPs and --indep-pairwise 50 10 0.5 filtering resulted in 41,120 SNPs), and a NJ tree using 500 bootstraps was created (https://itol.embl.de/tree/14915521019236421655393224).

### Reference-free clustering of blast fungus isolates based on K-mer sharing

To avoid potential biases due to differences in nucleotide divergence between the *M. oryzae* reference genome (rice blast fungus isolate 70-15) and each of the host-specific lineages, we leveraged the unmapped raw sequences of 80 representative *Magnaporthe* samples (**Table S2**) to estimate k-mer-based distances between isolates (https://github.com/smlatorreo/Magnaporthe-Germany). To eliminate possible adapter leftovers, we used AdapterRemoval (v2.3.1) [61] with default parameters. To calculate the k-mer-based pairwise distances we used Mash (v2.3) [68] with two different sets of parameters. The first consisted of sketch sizes of 100,000, k-mer sizes of 21 nucleotides, and a support of a minimum of 2 k-mer copies as a noise filter. The second settings consisted of sketch sizes of 1’000,000, k-mer sizes of 21 nucleotides, and a support of a minimum of 3 k-mer copies as a noise filter. Lastly, we created a pairwise distance matrix and clustered the isolates using the Neighbor joining algorithm from the R ape package [69]. The resulting NJ trees were visualized in iTol [67] (first parameter set: https://itol.embl.de/tree/82159717183891657273196, **Fig S2A** and second parameter set: https://itol.embl.de/tree/15818179229445911656070594, **Fig S2B**).

### Determination of mating type and candidate effector gene repertoire

To determine the mating type of the four sequenced German isolates, we searched for sequence similarity to the MAT1-1 and MAT1-2 mating type locus idiomorphs [31] using BLASTN. As a control, we also tested isolates whose mating type have been previously determined [32]. In addition, to compare the repertoire of candidate effector genes present in the German blast fungus isolates and identify presence/absence variation between them, we extracted the DNA coding sequences of a set of previously identified, either experimentally validated or in silico predicted effectors [35]. The candidate effectors were filtered to exclude highly similar (≥ 90% sequence identity) sequences, resulting in a total of 178 effectors [36]. These effectors originate from isolates infecting a variety of host grasses. We searched for sequence similarity between these predicted effectors across eight *M. oryzae* and eight *M. grisea* genomes using BLASTN [70]. An effector was deemed present if it showed ≥ 90% sequence identity and ≥ 90% sequence coverage. Effector presence/absence variation was plotted using the heatmap.2 function in gplots R package [71].

## Acknowledgements

We thank all members of the Kamoun lab, especially Daniel Lüdke and the Gretchens. We also thank the BLASTOFF team at the Sainsbury Laboratory for valuable discussions as well as Thomas Langner and Manfred Langner for their help with sampling and shipping of the samples.

## Data availability

The authors confirm that all data underlying the findings are fully available without restriction. Sequencing reads were deposited in the European Nucleotide Archive (ENA) under study accession number PRJEB53658 and run accessions ERR9866173 (GE12B), ERR9866254 (GE10A2), ERR9883755 and ERR9883756 (GE3) and ERR9866258 and ERR9866259 (GE16_2). In addition, the four whole-genome assemblies are available under GenBank accession numbers GCA_944989255.1 (GE12B), GCA_944989335.1 (GE10A2), GCA_944989265.1 (GE3) and GCA_944952825.1 (GE16_2). Detailed information is available in Barragan et al., (2022, Zenodo; https://zenodo.org/record/7010081#.YwkFw-zML0p).

## Author Contributions

**Conceptualization:** ACB, SK,TL.

**Formal analysis:** ACB, SML, PGM, AH, JW, AM,TL.

**Funding acquisition:** SK, HAB.

**Project administration:** SK,TL.

**Supervision:** HAB, SK,TL.

**Writing – original draft:** ACB.

**Writing – review & editing:** ACB, SML, HAB, SK,TL.

## References

1. Ristaino JB,Anderson PK, Bebber DP, Brauman KA, Cunniffe NJ, Fedoroff NV, et al. The persistent threat of emerging plant disease pandemics to global food security. Proc Natl Acad Sci U S A. 2021;118. doi: 10.1073/pnas.2022239118

2. Thines M.An evolutionary framework for host shifts - jumping ships for survival. New Phytol. 2019;224: 605–617.

3. Rahman MT, Sobur MA, Islam MS, Ievy S, Hossain MJ, El Zowalaty ME, et al. Zoonotic Diseases: Etiology, Impact, and Control. Microorganisms. 2020;8. doi:10.3390/microorganisms8091405

4. Carvajal-Yepes M, Cardwell K, Nelson A, Garrett KA, Giovani B, Saunders DGO, et al. A global surveillance system for crop diseases. Science. 2019;364: 1237–1239.

5. McCann HC. Skirmish or war: the emergence of agricultural plant pathogens. Curr Opin Plant Biol. 2020;56: 147–152.

6. Ou SH. Rice Diseases. IRRI; 1985.

7. Couch BC, Kohn LM.A multilocus gene genealogy concordant with host preference indicates segregation of a new species, Magnaporthe oryzae, from M. grisea. Mycologia. 2002;94: 683–693.

8. Klaubauf S,Tharreau D, Fournier E, Groenewald JZ, Crous PW, de Vries RP, et al. Resolving the polyphyletic nature of Pyricularia (Pyriculariaceae). Stud Mycol. 2014;79: 85–120.

9. Gladieux P, Ravel S, Rieux A, Cros-Arteil S, Adreit H, Milazzo J, et al. Coexistence of Multiple Endemic and Pandemic Lineages of the Rice Blast Pathogen. MBio. 2018;9. doi:10.1128/mBio.01806-17

10. Thierry M, Charriat F, Milazzo J,Adreit H, Ravel S, Cros-Arteil S, et al. Ecological Differentiation Among Globally Distributed Lineages of the Rice Blast Fungus Pyricularia oryzae. bioRxiv. 2021. p. 2020.06.02.129296. doi:10.1101/2020.06.02.129296

11. Couch BC, Fudal I, Lebrun M-H,Tharreau D,Valent B, van Kim P, et al. Origins of host-specific populations of the blast pathogen Magnaporthe oryzae in crop domestication with subsequent expansion of pandemic clones on rice and weeds of rice. Genetics. 2005;170: 613–630.

12. Inoue Y,Vy TTP,Yoshida K,Asano H, Mitsuoka C,Asuke S, et al. Evolution of the wheat blast fungus through functional losses in a host specificity determinant. Science. 2017;357: 80–83.

13. Pordel A, Ravel S, Charriat F, Gladieux P, Cros-Arteil S, Milazzo J, et al.Tracing the Origin and Evolutionary History of Pyricularia oryzae Infecting Maize and Barnyard Grass. Phytopathology. 2021;111: 128–136.

14. Malaker PK, Barma NCD,Tiwari TP, Collis WJ, Duveiller E, Singh PK, et al. First Report of Wheat Blast Caused by Magnaporthe oryzae Pathotype triticum in Bangladesh. Plant Dis. 2016;100: 2330–2330.

15. Islam MT,Tofazzal Islam M, Croll D, Gladieux P, Soanes DM, Persoons A, et al. Emergence of wheat blast in Bangladesh was caused by a South American lineage of Magnaporthe oryzae. BMC Biology. 2016. doi: 10.1186/s12915-016-0309-7

16. Tembo B, Mulenga RM, Sichilima S, M’siska KK, Mwale M, Chikoti PC, et al. Detection and characterization of fungus (Magnaporthe oryzae pathotype Triticum) causing wheat blast disease on rain-fed grown wheat (Triticum aestivum L.) in Zambia. PLoS One. 2020;15: e0238724.

17. Roumen E, Levy M, Notteghem JL. Characterisation of the European pathogen population of Magnaporthe grisea by DNA fingerprinting and pathotype analysis. Eur J Plant Pathol. 1997;103: 363–371.

18. Beck HE, Zimmermann NE, McVicar TR,Vergopolan N, Berg A,Wood EF. Present and future Köppen-Geiger climate classification maps at 1-km resolution. Sci Data. 2018;5: 180214.

19. Bebber DP, Ramotowski MAT, Gurr SJ. Crop pests and pathogens move polewards in a warming world. Nat Clim Chang. 2013;3: 985–988.

20. Nnadi NE, Carter DA. Climate change and the emergence of fungal pathogens. PLoS Pathog. 2021;17: e1009503.

21. Goertz A, Zuehlke S, Spiteller M, Steiner U, Dehne HW,Waalwijk C, et al. Fusarium species and mycotoxin profiles on commercial maize hybrids in Germany. Eur J Plant Pathol. 2010;128: 101–111.

22. Barragan AC,Win J, Harant A, Kamoun S, Langner T. Genome assemblies of the blast fungus Magnaporthe (Syn. Pyricularia) spp. collected from wild grasses in Germany. Zenodo; 2022. doi: 10.5281/ZENODO.7010081

23. Dean RA,Talbot NJ, Ebbole DJ, Farman ML, Mitchell TK, Orbach MJ, et al.The genome sequence of the rice blast fungus Magnaporthe grisea. Nature. 2005;434: 980–986.

24. Yoshida K, Saunders DGO, Mitsuoka C, Natsume S, Kosugi S, Saitoh H, et al. Host specialization of the blast fungus Magnaporthe oryzae is associated with dynamic gain and loss of genes linked to transposable elements. BMC Genomics. 2016;17: 370.

25. Gladieux P, Condon B, Ravel S, Soanes D, Maciel JLN, Nhani A Jr, et al. Gene Flow between Divergent Cereal-and Grass-Specific Lineages of the Rice Blast Fungus Magnaporthe oryzae. MBio. 2018;9. doi: 10.1128/mBio.01219-17

26. Islam MT, Kim K-H, Choi J.Wheat Blast in Bangladesh: The Current Situation and Future Impacts. Plant Pathol J. 2019;35: 1–10.

27. Peng Z, Oliveira-Garcia E, Lin G, Hu Y, Dalby M, Migeon P, et al. Effector gene reshuffling involves dispensable mini-chromosomes in the wheat blast fungus. PLoS Genet. 2019;15: e1008272.

28. Latorre SM,Were VM, Foster AJ, Langner T, Malmgren A, Harant A, et al.A pandemic clonal lineage of the wheat blast fungus. 2022. doi: 10.1101/2022.06.06.494979

29. Liu S, Lin G, Ramachandran SR, Cruppe G, Cook D, Pedley KF, et al. Rapid mini-chromosome divergence among fungal isolates causing wheat blast outbreaks in Bangladesh and Zambia. bioRxiv. 2022. p. 2022.06.18.496690. doi:10.1101/2022.06.18.496690

30. Wilson AM,Wilken PM, van der Nest MA,Wingfield MJ,Wingfield BD. It’s All in the Genes:The Regulatory Pathways of Sexual Reproduction in Filamentous Ascomycetes. Genes. 2019;10. doi: 10.3390/genes10050330

31. Kang S, Chumley FG,Valent B. Isolation of the mating-type genes of the phytopathogenic fungus Magnaporthe grisea using genomic subtraction. Genetics. 1994;138: 289–296.

32. Latorre SM,Were VM, Langer T, Foster AJ,Win J, Kamoun S, et al.A curated set of mating-type assignment for the blast fungus (Magnaporthales). Zenodo; 2022. doi:10.5281/ZENODO.6369833

33. Kang S, Lebrun MH, Farrall L,Valent B. Gain of virulence caused by insertion of a Pot3 transposon in a Magnaporthe grisea avirulence gene. Mol Plant Microbe Interact. 2001;14: 671–674.

34. Chiapello H, Mallet L, Guérin C,Aguileta G,Amselem J, Kroj T, et al. Deciphering Genome Content and Evolutionary Relationships of Isolates from the Fungus Magnaporthe oryzae Attacking Different Host Plants. Genome Biol Evol. 2015;7: 2896–2912.

35. Petit-Houdenot Y, Langner T, Harant A,Win J, Kamoun S.A Clone Resource of Magnaporthe oryzae Effectors That Share Sequence and Structural Similarities Across Host-Specific Lineages. Mol Plant Microbe Interact. 2020;33: 1032–1035.

36. Latorre SM, Reyes-Avila CS, Malmgren A,Win J, Kamoun S, Burbano HA. Differential loss of effector genes in three recently expanded pandemic clonal lineages of the rice blast fungus. BMC Biol. 2020;18: 88.

37. de Guillen K, Ortiz-Vallejo D, Gracy J, Fournier E, Kroj T, Padilla A. Structure Analysis Uncovers a Highly Diverse but Structurally Conserved Effector Family in Phytopathogenic Fungi. PLoS Pathog. 2015;11: e1005228.

38. Kim K-T, Ko J, Song H, Choi G, Kim H, Jeon J, et al. Evolution of the Genes Encoding Effector Candidates Within Multiple Pathotypes of Magnaporthe oryzae. Front Microbiol. 2019;10: 2575.

39. Chuma I, Isobe C, Hotta Y, Ibaragi K, Futamata N, Kusaba M, et al. Multiple translocation of the AVR-Pita effector gene among chromosomes of the rice blast fungus Magnaporthe oryzae and related species. PLoS Pathog. 2011;7: e1002147.

40. Cesari S,Thilliez G, Ribot C, Chalvon V, Michel C, Jauneau A, et al.The rice resistance protein pair RGA4/RGA5 recognizes the Magnaporthe oryzae effectors AVR-Pia and AVR1-CO39 by direct binding. Plant Cell. 2013;25: 1463–1481.

41. Leff B, Ramankutty N, Foley JA. Geographic distribution of major crops across the world. Global Biogeochem Cycles. 2004;18. doi: 10.1029/2003gb002108

42. Morris CE, Moury B. Revisiting the Concept of Host Range of Plant Pathogens.Annu Rev Phytopathol. 2019;57: 63–90.

43. Farman M, Peterson G, Chen L, Starnes J,Valent B, Bachi P, et al.The Lolium Pathotype of Magnaporthe oryzae Recovered from a Single Blasted Wheat Plant in the United States. Plant Dis. 2017;101: 684–692.

44. Hyon G-S, Nga NTT, Chuma I, Inoue Y, Asano H, Murata N, et al. Characterization of interactions between barley and various host-specific subgroups of Magnaporthe oryzae and M. grisea. J Gen Plant Pathol. 2012;78: 237–246.

45. Chung H, Goh J, Han S-S, Roh J-H, Kim Y, Heu S, et al. Comparative Pathogenicity and Host Ranges of Magnaporthe oryzae and Related Species. Plant Pathol J. 2020;36: 305–313.

46. Saleh D, Xu P, Shen Y, Li C,Adreit H, Milazzo J, et al. Sex at the origin: an Asian population of the rice blast fungus Magnaporthe oryzae reproduces sexually. Mol Ecol. 2012;21: 1330–1344.

47. Thierry M, Charriat F, Milazzo J,Adreit H, Ravel S, Cros-Arteil S, et al. Maintenance of divergent lineages of the Rice Blast Fungus Pyricularia oryzae through niche separation, loss of sex and post-mating genetic incompatibilities. PLoS Pathog. 2022;18: e1010687.

48. Sweigard JA, Carroll AM, Kang S, Farrall L, Chumley FG,Valent B. Identification, cloning, and characterization of PWL2, a gene for host species specificity in the rice blast fungus. Plant Cell. 1995;7: 1221–1233.

49. Nakayama H, Nagamine T, Hayashi N. Genetic Variation of Blast Resistance in Foxtail Millet (Setaria italica (L.) P. Beauv.) and its Geographic Distribution. Genet Resour Crop Evol. 2005;52: 863–868.

50. Andersen EJ, Nepal MP. Genetic diversity of disease resistance genes in foxtail millet (Setaria italica L.). Plant Gene. 2017;10: 8–16.

51. Shimizu M, Hirabuchi A, Sugihara Y, Abe A,Takeda T, Kobayashi M, et al.A genetically linked pair of NLR immune receptors show contrasting patterns of evolution. bioRxiv. 2021. p. 2021.09.01.458560. doi: 10.1101/2021.09.01.458560

52. Hersbach H, Bell B, Berrisford P, Biavati G, Horányi A, Muñoz Sabater J, et al. ERA5 monthly averaged data on single levels from 1979 to present. Copernicus Climate Change Service (C3S) Climate Data Store (CDS). 2019;10: 252–266.

53. Meinshausen M, Nicholls ZRJ, Lewis J, Gidden MJ, Vogel E, Freund M, et al.The shared socio-economic pathway (SSP) greenhouse gas concentrations and their extensions to 2500. Geosci Model Dev. 2020;13: 3571–3605.

54. Ades M,Adler R,Allan R,Allan RP, Anderson J,Argüez A, et al. Global Climate. Bull Am Meteorol Soc. 2020;101: S9–S128.

55. Tebaldi C, Debeire K, Eyring V, Fischer E, Fyfe J, Friedlingstein P, et al. Climate model projections from the Scenario Model Intercomparison Project (ScenarioMIP) of CMIP6. Earth Syst Dyn. 2021;12: 253–293.

56. Monfreda C, Ramankutty N, Foley JA. Farming the planet: 2. Geographic distribution of crop areas, yields, physiological types, and net primary production in the year 2000. Global Biogeochem Cycles. 2008;22. doi: 10.1029/2007gb002947

57. Walter Kahlenborn, Luise Porst, Maike Voß, Uta Fritsch, Kathrin Renner, Marc Zebisch, Mareike Wolf, Konstanze Schönthaler, Inke Schauser. Klimawirkungs- und Risikoanalyse für Deutschland 2021 (Kurzfassung). 2021.

58. Kahle D,Wickham H. ggmap: Spatial Visualization with ggplot2. R J. 2013;5: 144–161.

59. Dubina EV, Alabushev AV, Kostylev PI, Kharchenko ES, Ruban MG,Aniskina YV, et al. Biodiversity of Pyricularia oryzae Cav. in rice-growing regions of the south of Russia using PCR method. Physiol Mol Biol Plants. 2020;26: 289–303.

60. Chung H, Jeong DG, Lee J-H, Kang IJ, Shim H-K, An CJ, et al. Outbreak of Rice Blast Disease at Yeoju of Korea in 2020. Plant Pathol J. 2022;38: 46–51.

61. Schubert M, Lindgreen S, Orlando L.AdapterRemoval v2: rapid adapter trimming, identification, and read merging. BMC Res Notes. 2016;9: 88.

62. Li H.Aligning sequence reads, clone sequences and assembly contigs with BWA-MEM. arXiv [q-bio.GN]. 2013.Available: http://arxiv.org/abs/1303.3997

63. Danecek P, Bonfield JK, Liddle J, Marshall J, Ohan V, Pollard MO, et al.Twelve years of SAMtools and BCFtools. Gigascience. 2021;10. doi: 10.1093/gigascience/giab008

64. McKenna A, Hanna M, Banks E, Sivachenko A, Cibulskis K, Kernytsky A, et al.The Genome Analysis Toolkit: a MapReduce framework for analyzing next-generation DNA sequencing data. Genome Res. 2010;20: 1297–1303.

65. Latorre SM, Langner T, Malmgren A,Win J, Kamoun S, Burbano HA. SNP calling parameters have minimal impact on population structure and divergence time estimates for the rice blast fungus. bioRxiv. 2022. p. 2022.03.06.482794. doi:10.1101/2022.03.06.482794

66. Kumar S, Stecher G, Li M, Knyaz C,Tamura K. MEGA X: Molecular Evolutionary Genetics Analysis across Computing Platforms. Mol Biol Evol. 2018;35: 1547–1549.

67. Letunic I, Bork P. Interactive Tree Of Life (iTOL) v5: an online tool for phylogenetic tree display and annotation. Nucleic Acids Res. 2021;49:W293–W296.

68. Ondov BD,Treangen TJ, Melsted P, Mallonee AB, Bergman NH, Koren S, et al Mash: fast genome and metagenome distance estimation using MinHash. Genome Biol. 2016;17: 132.

69. Paradis E, Schliep K. ape 5.0: an environment for modern phylogenetics and evolutionary analyses in R. Bioinformatics. 2019;35: 526–528.

70. Altschul SF, Gish W, Miller W, Myers EW, Lipman DJ. Basic local alignment search tool. J Mol Biol. 1990;215: 403–410.

71. Warnes GR, Bolker B, Bonebakker L, Gentleman R, Huber W, Liaw A, et al. gplots: Various R programming tools for plotting data. R package version. 2009;2: 1.

