## Supplemental_Figures for "Wild grass isolates of *Magnaporthe* (Syn. *Pyricularia*) spp. from Germany can cause blast disease on cereal crops"

**Fig S1.** SNP-based Neighbor Joining trees indicate that *M. oryzae* and its sister species *M. grisea* are present in Germany. Related to Fig 2.

**Fig S2.** K-mer based clustering indicate that *M. oryzae* and its sister species *M. grisea* are present in Germany. Related to Fig 2.

**Fig S3.** German *Setaria* spp.-infecting *M. oryzae* isolates can complete their infection cycle on barley and wheat cultivars. Related to Fig 3.

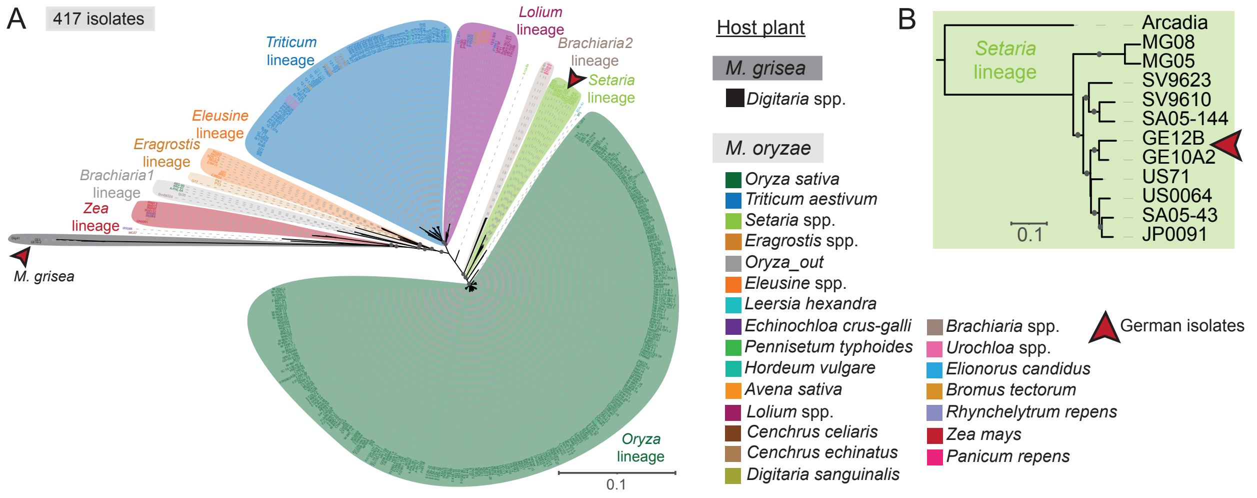


**Fig S1. SNP-based Neighbor Joining trees indicate that *M. oryzae* and its sister species *M. grisea* are present in Germany. A.** SNP-based genome-wide NJ tree of 417 *Magnaporthe* isolates color-coded by host plant and blast fungus lineage. **B.** SNP-based genome-wide NJ tree of members of the *Setaria*-infecting *M. oryzae* lineage. Red arrows highlight the German samples. German *M. oryzae* isolates are genetically most similar to each other. *Digitaria* spp.-infecting German samples (GE3 and GE16_2) cluster with the *M. grisea* sample Dig41 (black), while the *Setaria* spp.-infecting German samples (GE12B and GE10A2, red) cluster with *Setaria* spp.-infecting *M. oryzae* isolates (light green). Relevant bootstrapping values ≥ 0.9 are shown as gray circles. Scale bar indicates nucleotide substitutions per position. Related to Fig 2.

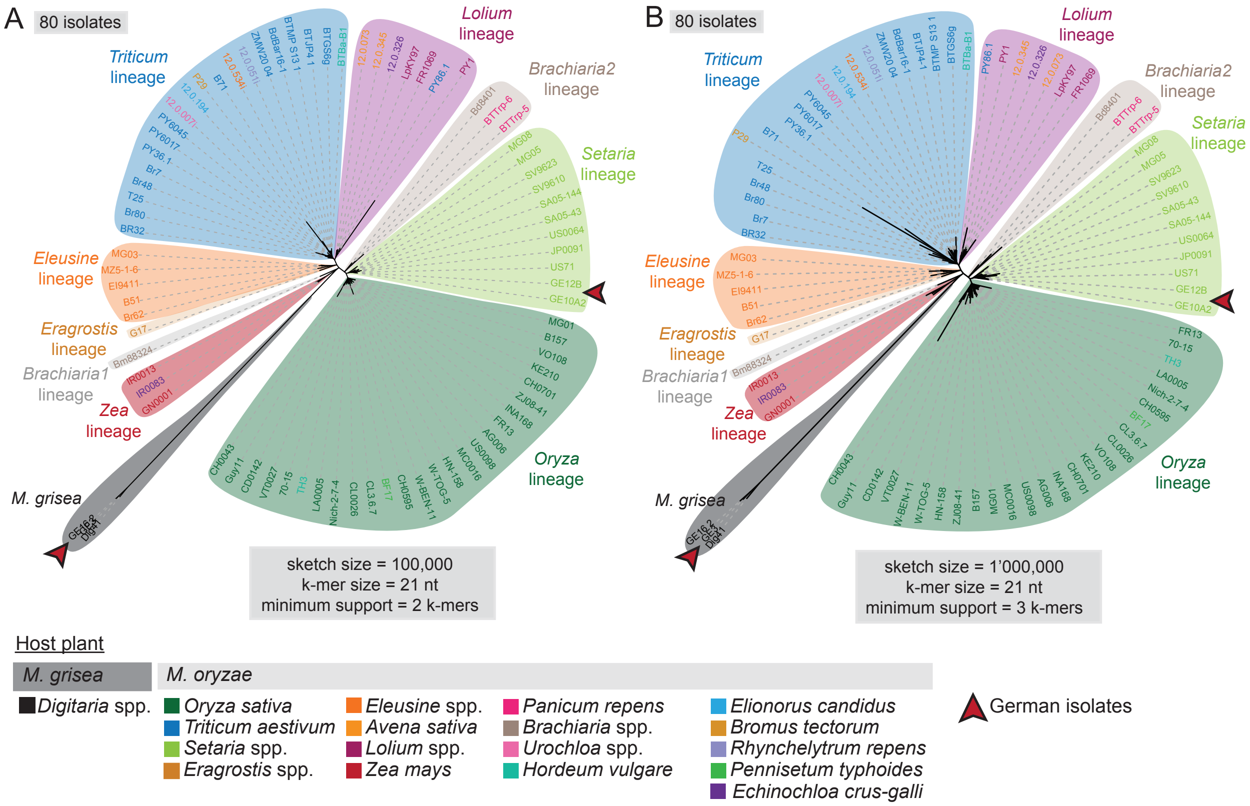


**Fig S2. K-mer based clustering indicate that *M. oryzae* and its sister species *M. grisea* are present in Germany.** Reference-free clustering of 80 blast fungus isolates (Table S2) based on k-mer sharing. Red arrows highlight the German samples. Two different parameter settings were chosen (see Methods). In both cases *Digitaria* spp.-infecting German samples (GE3 and GE16_2) cluster with the *M. grisea* sample Dig41 (black), while the *Setaria* spp.-infecting German samples (GE12B and GE10A2, red) cluster with *Setaria* spp.-infecting *M. oryzae* isolates (light green). Related to Fig 2.

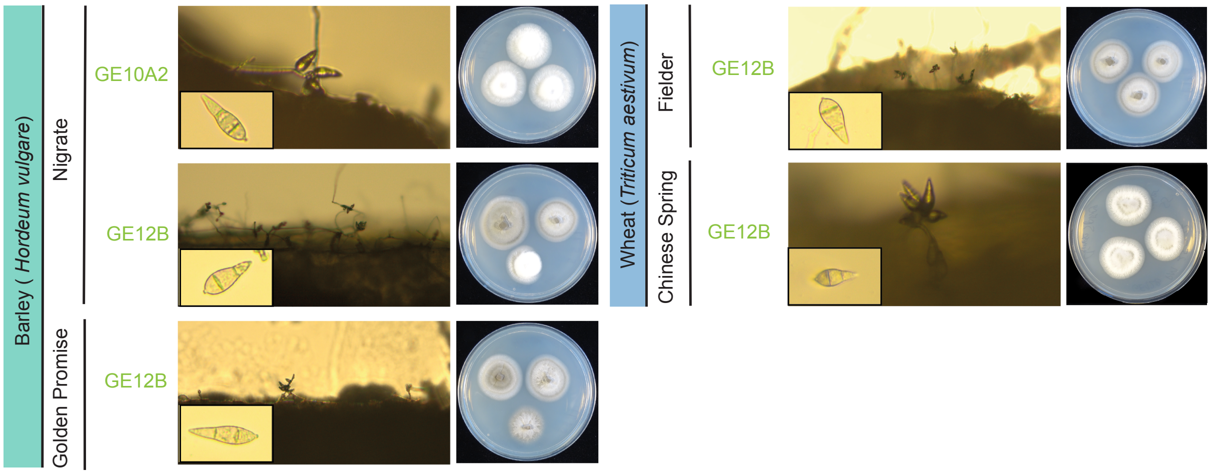


**Fig S3. German *Setaria* spp.-infecting *M. oryzae* isolates can complete their infection cycle on barley and wheat cultivars.** Formation of conidiophores on barley and wheat cultivars. Lesions from drop infection assays were transferred to 2% water agar and formation of conidiophores was observed microscopically after 24 hours. Conidiophore formation was observed for all tested cultivars from infections with the *M. oryzae* isolate GE12B and from infections of the barley cultivar Nigrate with *M. oryzae* isolate GE10A2. Conidia produced on these host plants were viable. Viability of the spores was tested by cultivating single spores on complete medium, which led to the formation of typical blast fungus colonies. Related to Fig 3.
